# High-resolution cryo-EM structure of photosystem II: Effects of electron beam damage

**DOI:** 10.1101/2020.10.18.344648

**Authors:** Koji Kato, Naoyuki Miyazaki, Tasuku Hamaguchi, Yoshiki Nakajima, Fusamichi Akita, Koji Yonekura, Jian-Ren Shen

## Abstract

Photosystem II (PSII) plays a key role in water-splitting and oxygen evolution. X-ray crystallography has revealed its atomic structure and some intermediate structures. However, these structures are in the crystalline state, and its final state structure has not been solved because of the low efficiencies of the S-state transitions in the crystals. Here we analyzed the structure of PSII in solution at 1.95 Å resolution by single-particle cryo-electron microscopy (cryo-EM). The structure obtained is similar to the crystal structure, but a PsbY subunit was visible in the cryo-EM structure, indicating that it represents its physiological state more closely. Electron beam damage was observed at a high-dose in the regions that were easily affected by redox states, which was reduced by reducing the electron dose. This study will serve as a good indicator for determining damage-free cryo-EM structures of not only PSII but also all biological samples, especially redox-active metalloproteins.

## Introduction

PSII is a multi-subunit pigment-protein complex embedded in the thylakoid membranes of higher plants, green algae and cyanobacteria, and is the only molecular machine capable of oxidizing water by use of visible light in the nature. Water molecules are split into electrons, hydrogen atoms and oxygen molecules at the catalytic center of PSII, namely, the oxygen-evolving complex (OEC), through four electron and/or proton removing steps as described in the Si-state cycle (with i = 0–4, where i indicates the number of oxidative equivalents accumulated)^1^.

In order to elucidate the mechanism of the water-splitting reaction, the structure of PSII has been studied extensively by X-ray diffraction (XRD) with synchrotron radiation (SR)^2–6^. The SR structure of PSII at an atomic resolution revealed that OEC is a Mn_4_CaO_5_ cluster organized into a distorted-chair form, in which the cuboidal part is composed of Mn_3_CaO_4_ and the outer manganese is attached to the cuboid via two μ-oxo-bridges^6^. However, based on the extended X-ray absorption fluorescence spectra (EXAFS) analysis, the dose used for collecting the SR structure at 1.9 Å resolution may cause 25% of the Mn ions in OEC to be reduced to 2^+^ ions, causing some elongations in the Mn-Mn distances in the structure^7^. This issue is overcome by the use of X-ray free electron lasers (XFEL), which provide X-ray pulses with ultra-short durations that enable collection of the diffraction data before onset of the radiation damage (diffraction before destruction)^8^. Using XFELs, radiation damage free structure of PSII was solved at a high resolution by an approach called fixed-target serial rotational crystallography, which uses multiple large PSII crystals by a shot-and-move/rotation method^9,10^. The result showed a shortening of 0.1-0.2 Å in some of the Mn-Mn distances, indicating that the structure represents a damage free one^10^. By a combination of serial femtosecond X-ray crystallography (SFX) with XFELs and small crystals, structures of S-state intermediates up to S_3_-state were analyzed by pump-probe experiments where snapshot diffraction images were collected from flash-illuminated PSII crystals^11–14^. These results demonstrated the appearance of a new oxygen atom O6 (Ox) close to O5 between Mn1 and Mn4 upon two flashes, suggesting insertion of a water molecule in the S_2_ → S_3_ transition for O=O bond formation. However, all these studies were conducted with PSII crystals, and the efficiencies of the S-state transitions in the microcrystals were reported to be slightly lower compared with those in solution using light-induced Fourier transform infrared difference spectroscopy^15^. Moreover, it is unknown if the structure of PSII in the crystalline state is the same as those in the solution.

Cryo-electron microscopy (cryo-EM) can solve the structures of proteins in solution without crystallization, which may represent the physiological states of proteins more closely. It can also analyze the dynamic changes of proteins in solutions in the time range of ms, provided that cooling of the samples is rapid enough. In recent years, the technique of cryo-EM has been developed rapidly, and the resolutions of structures that can be solved by cryo-EM are increased dramatically^16–18^. However, there is also the issue of damage caused by the electron beam during cryo-EM data collection, even though the cryo-EM is usually conducted at a low temperature. Radiation damage has been extensively studied with X-rays, and it has been shown that the damage mainly manifests as breakage of disulfide bonds, decarboxylation of acidic amino and photoreduction of metal centers^7,19–21^. The damage caused by electron beams have also been shown in cryo-EM analysis^17^. In order to obtain a high resolution, however, cryo-EM studies are usually conducted at a high-dose of electron beams without paying much attention to the electron beam damage. In this paper, we analyzed the structure of PSII in ice by cryo-EM at a resolution of 1.95 Å, and investigated the electron beam damage to PSII, especially its OEC, upon dose accumulation. We show that the structure of PSII analyzed by cro-EM may represent the physiological state more closely, as it retains the PsbY subunit. However, it suffers from a severe electron beam damage at a high-dose, and this damage was reduced at a much decreased dose without a significant loss of resolution. These results are not only important for the analysis of the PSII structure in solution, but also provide important implications for all cryo-EM studies that use considerably high-doses for imaging.

## Result

### High resolution single particle analysis of the PSII

To obtain the high resolution structure of PSII, three data sets of single-particle images of the PSII dimer from *T. vulcanus* were collected using Thermo Fisher Scientific Titan Krios and JEOL CRYO ARM 300 at different conditions as summarized in Table 1. Because the sample for the 75 x k magnification using Titan contained 5% glycerol in the buffer, the sample was diluted ten times with the buffer without glycerol. The other samples did not contain the glycerol and were concentrated by PEG 1450 precipitation in the final step. Image processing yielded final resolutions of 2.22 Å for the data set collected at 75 x k magnification using Titan Krios (Titan-75k), 2.20 Å for the data set collected at 96 x k magnification using Titan Krios (Titan-96k), and 1.95 Å for the data set collected at 60 x k magnification using CRYO ARM 300 (ARM-60k) (Table 1, Table 2 and Supplementary Fig. 1-4). These results indicate that the quality of the cryo-EM density maps achieved were at the level comparable to those obtained with SR and XFEL previously^6,10^. The resolutions of the Titan-75k and Titan-96k data were almost the same, in spite of the different magnifications and buffers used. The resolution of the ARM-60k data was significantly better than that of the Titan-96k data, despite that the same buffer condition was used for the two data sets. The resolution achieved by cryo-EM depends on a number of factors, including sample quality, the type and preparations of cryo-grids used, the thickness of ice in the samples, microscope alignment and imaging conditions, etc. However, the major one could be the electron beam source. The CRYO ARM 300 microscope has a cold field emission gun (CFEG) that produces an electron beam with a high temporal-coherence and superior high-resolution signals^22^ over that from the Schottky emission gun equipped in the Krios microscope.

**Table 1.**
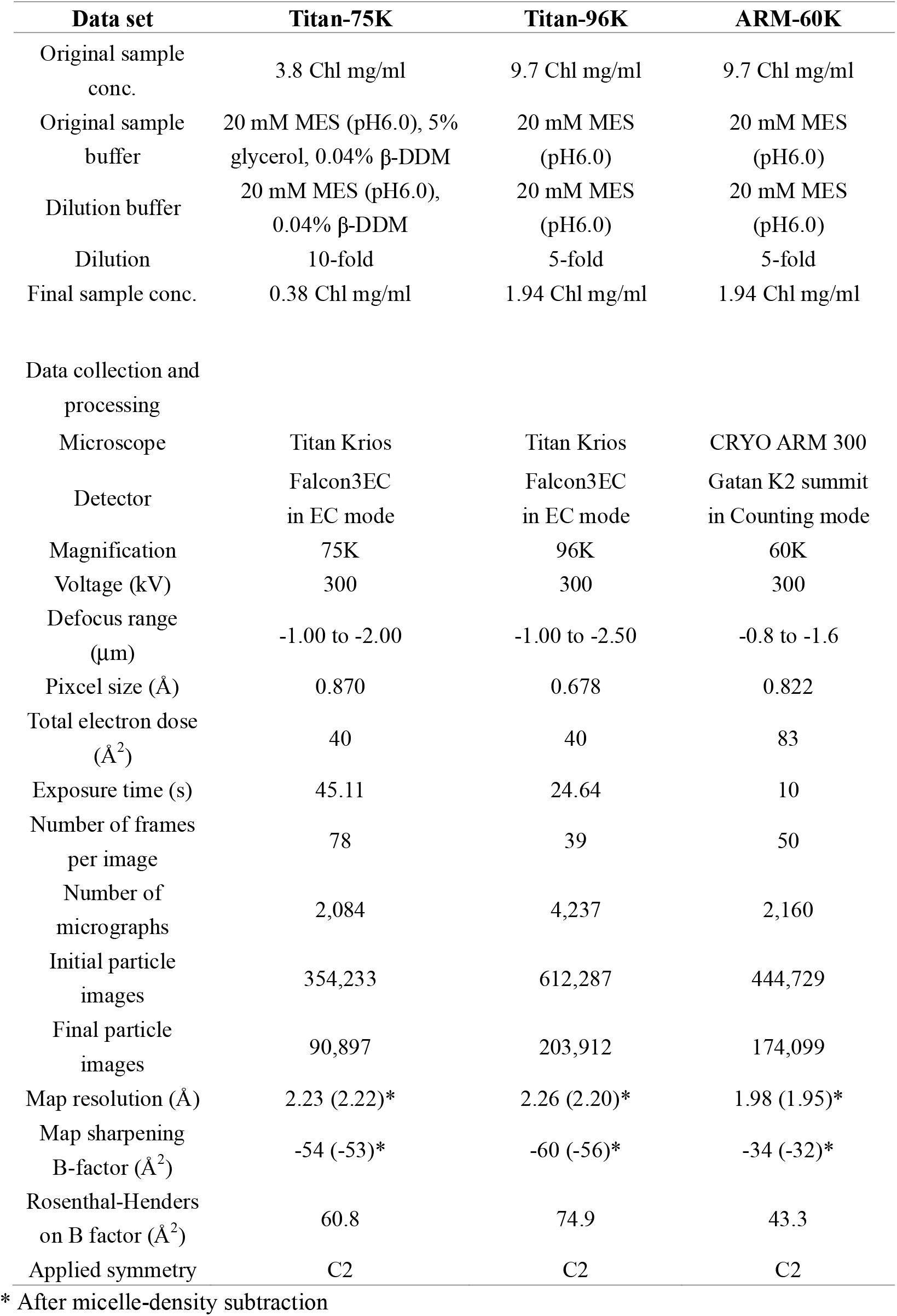
Sample preparation and Cryo-EM data collection parameters.

**Table 2.**
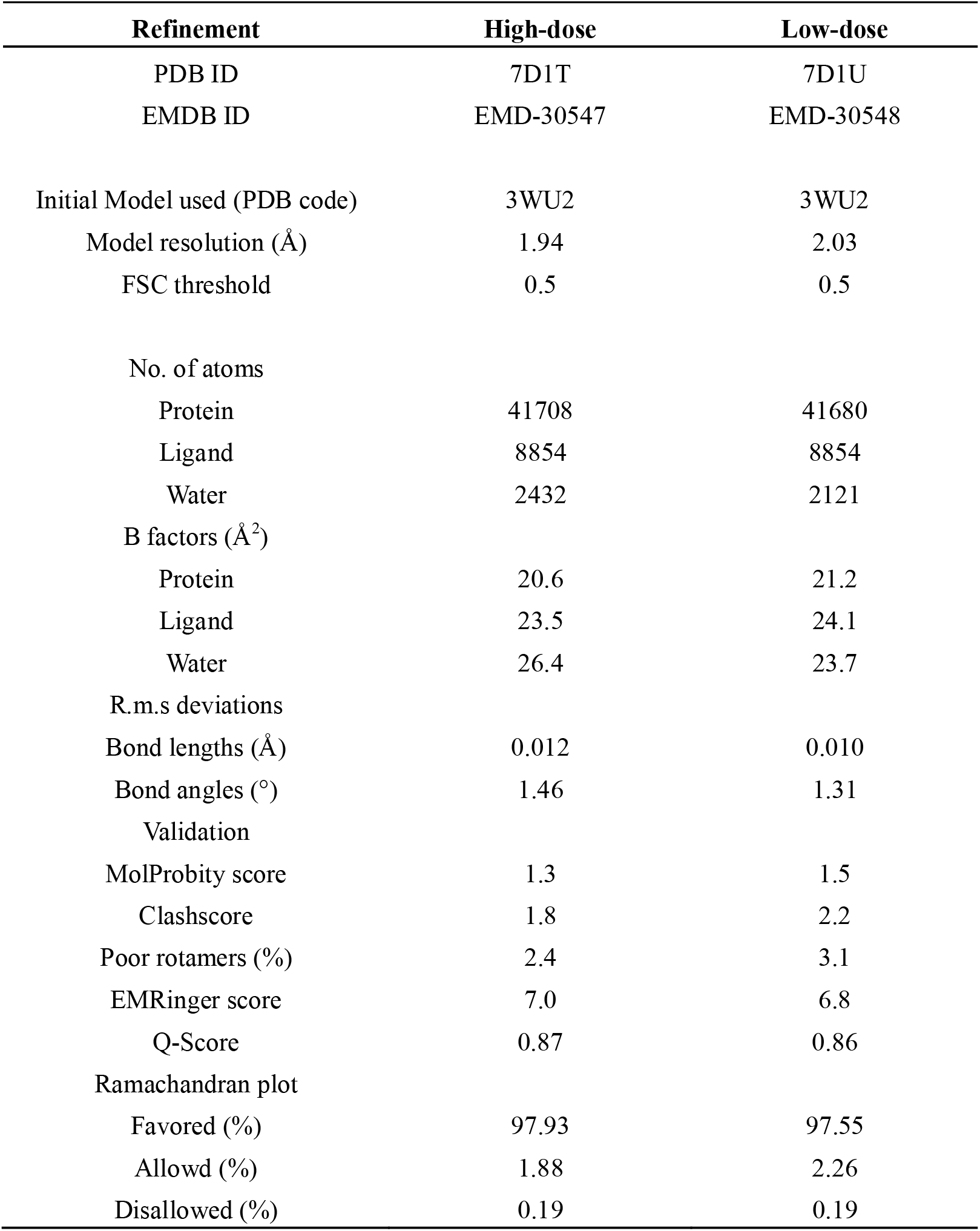
Statistics of data collection, processing and refinement.

| **Refinement** | **High-dose** | **Low-dose** |
| --- | --- | --- |
| PDB ID | 7D1T | 7D1U |
| EMDB ID | EMD-30547 | EMD-30548 |
| Initial Model used (PDB code) | 3WU2 | 3WU2 |
| Model resolution (Å) | 1.94 | 2.03 |
| FSC threshold | 0.5 | 0.5 |
| No. of atoms |  |  |
| Protein | 41708 | 41680 |
| Ligand | 8854 | 8854 |
| Water | 2432 | 2121 |
| B factors (Å<sup>2</sup>) |  |  |
| Protein | 20.6 | 21.2 |
| Ligand | 23.5 | 24.1 |
| Water | 26.4 | 23.7 |
| R.m.s deviations |  |  |
| Bond lengths (Å) | 0.012 | 0.010 |
| Bond angles (°) | 1.46 | 1.31 |
| Validation |  |  |
| MolProbity score | 1.3 | 1.5 |
| Clashscore | 1.8 | 2.2 |
| Poor rotamers (%) | 2.4 | 3.1 |
| EMRinger score | 7.0 | 6.8 |
| Q-Score | 0.87 | 0.86 |
| Ramachandran plot |  |  |
| Favored (%) | 97.93 | 97.55 |
| Allowd (%) | 1.88 | 2.26 |
| Disallowed (%) | 0.19 | 0.19 |

In Fig. 1, the squared inverse resolution of reconstructions achieved from random subsets of particles is plotted against the subset size on a logarithmic scale. This is known as Rosenthal-Henderson plot^23^. These plots indicated that the resolution is proportional to the log of particle size. The B-factors estimated from these plots are 60.8 for the Titan-75k data set, 74.9 for the Titan-96k data set and 43.3 Å^2^ for the ARM-60k data set. The ARM-60k data set again shows the lowest value, in agreement with its highest resolution.

**Fig. 1.**
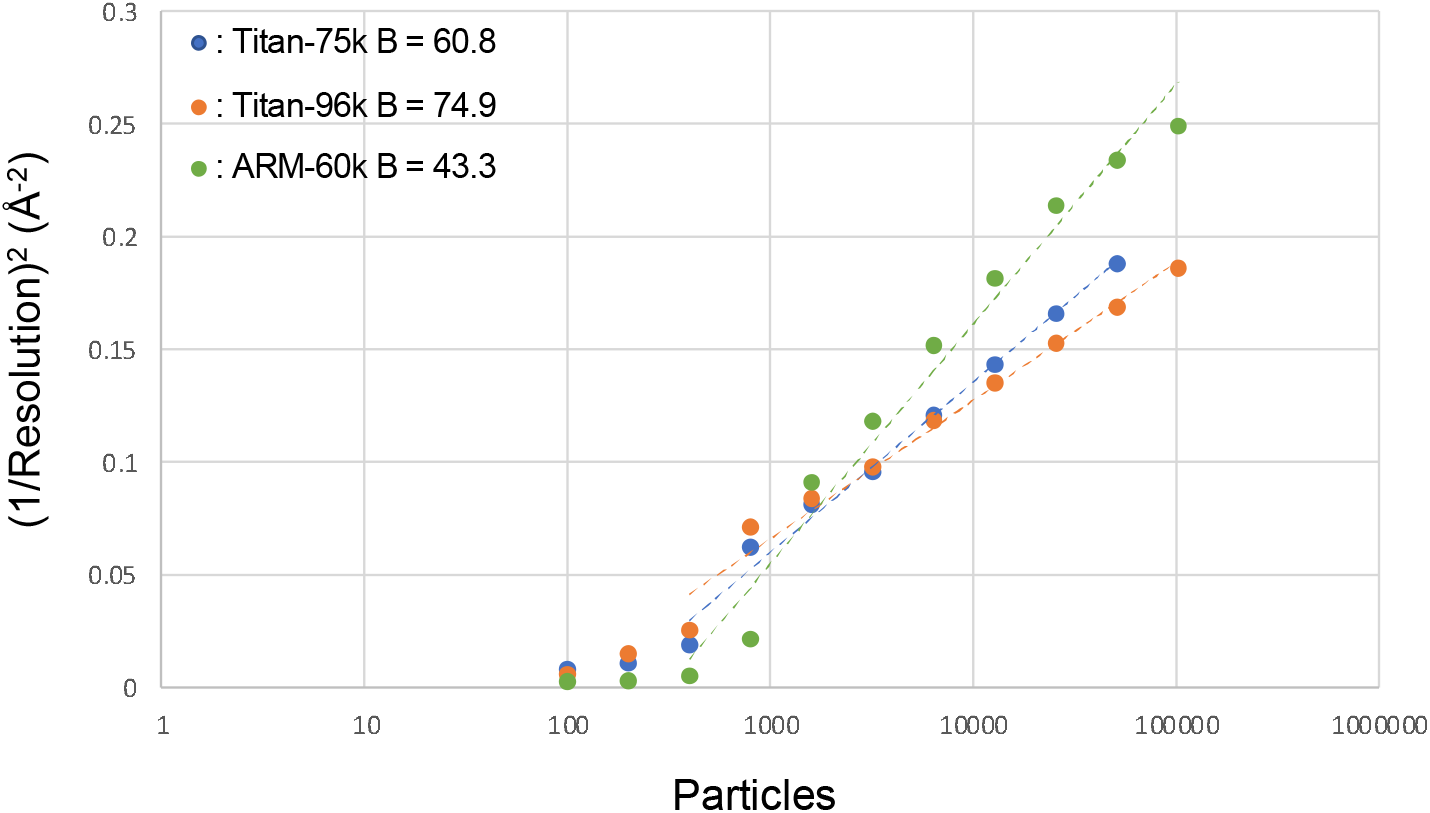
B-factor plot for the data sets of Titan-75k, Titan-96k and ARM-60k. B factor plot for the Titan-75k data set at a dose of 40 e^−^ Å^−2^ (blue), the Titan-96k data set at a dose of 40 e^−^ Å^−2^ (orange), and the ARM-60k data set at a dose of 83 e^−^ Å^−2^ (green).

### Overall structure of PSII

The overall atomic model of PSII was built based on the highest 1.95-Å resolution density map reconstructed from the ARM-60k data set. At this resolution, the features of cofactors and water molecules can be easily identified in the map (Fig. 2). The overall architecture of the PSII dimer from *T. vulcanus* is very similar to that of SR (PDB: 3WU2)^6^ and XFEL structures (PDB: 4UB6 and 4UB8)^10^, except for PsbY. The density of PsbY is present in one of the two monomers in the native (PDB: 4UB6)^10^ and the Sr^2+^-substituted PSII dimer structures (PDB: 4IL6)^24^ but absent in the SR structure (PDB: 3WU2)^6^. However, this density was seen in both sides of the PSII dimer in the cryo-EM structure, although the density is somewhat poorer compared with that of the other assigned subunits (Fig. 2a, f). This suggests that the cryo-EM structure more closely represents the native state of PSII.

**Fig. 2.**
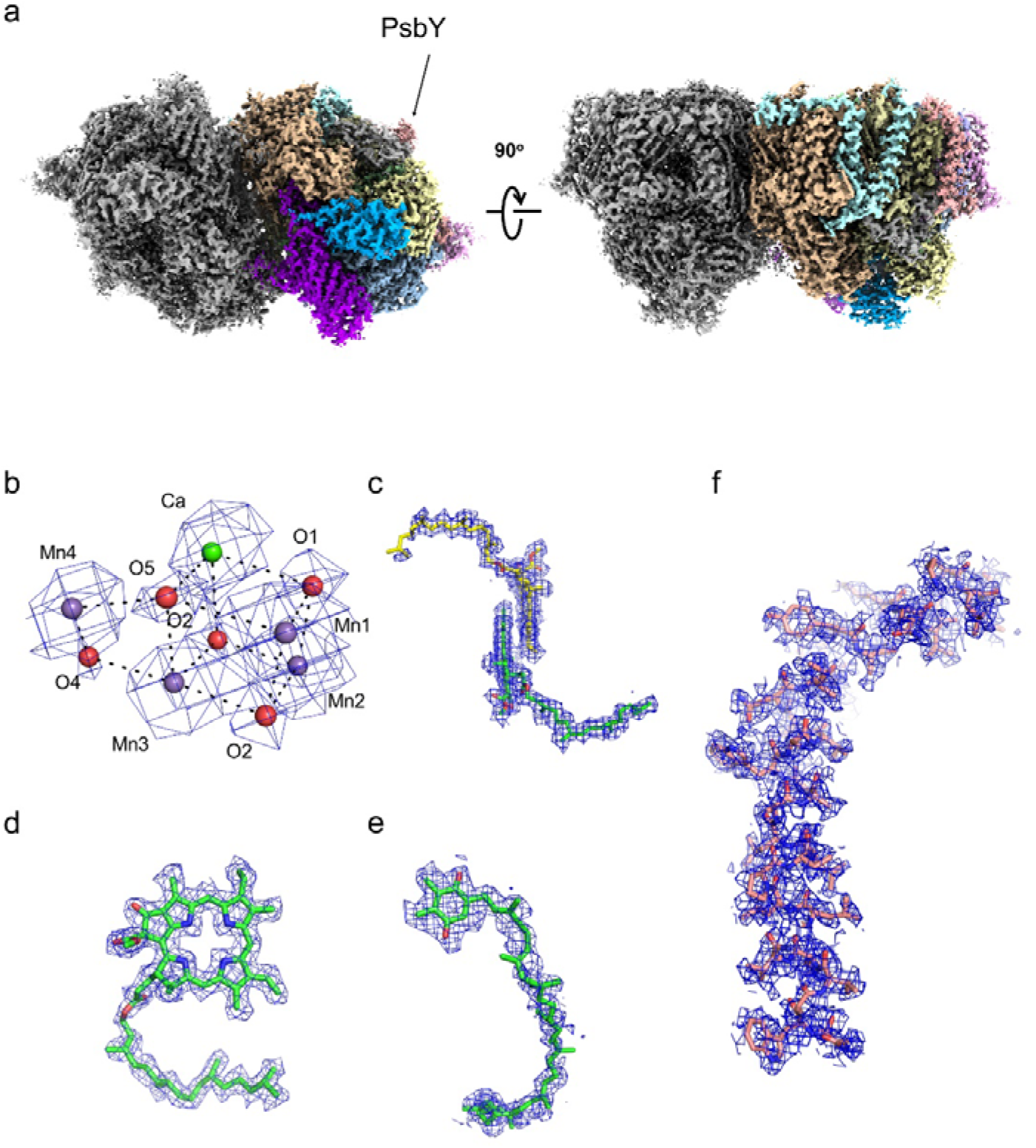
Overall structure of PSII at a high-dose. **a** The cryo-EM density of the PSII at 1.95 Å resolution from the ARM-60k data set. **b**-**e** The cryo-EM density of cofactors, OEC (**b**), P680 (**c**), pheophytin (**d**) and plastoquinone (Q_B_) (**e**), superposed with the refined model. **f** The density of PsbY superposed with the refined model. The densities were depicted at 5 σ.

The root mean square deviation (RMSD) is 0.40 Å for 5227 C_α_ atoms between the structures of cryo-EM and SR, and 0.46 Å for 5267 C_α_ atoms between the structures of cryo-EM and XFEL. Because the RMSD was 0.32 Å for 5241 C_α_ atoms between the SR and XFEL structures of PSII dimers, the cryo-EM structure is almost identical to the SR and XFEL structures at the backbone level. In the cryo-EM density map, we assigned 2432 water molecules at a contour level of 5 σ, which are slightly less than the number of waters assigned in the SR and XFEL structures^6,10^. The atomic displacement parameter (ADP) of the cryo-EM structure refined with Refmac5 in reciprocal space correlated well with that of the SR structure (Supplementary Fig. 5), although it may be somewhat overestimated in the cryo-EM structure. Since the cryo-EM map was subjected to B-factor sharpening with Postprocessing, it is not suitable to compare ADP values directly between cryo-EM and X-ray structures. Nevertheless, the relative ADP of the atoms in the molecule appears to reflect the characteristics of the map. The average ADP for the OEC atoms (13.8 Å^2^) were found to be lower than that observed in the overall protein atoms of the cryo-EM structure (20.6 Å^2^), suggesting that the structure of OEC was determined more reliably than that of the overall structure. This may be due to the presence of metal ions in the Mn_4_CaO_5_ cluster, which gives rise to higher cryo-EM density than that of lighter protein atoms.

### Electron beam damage to the PSII structure

Several regions of PSII were found to have different structures between cryo-EM and XFEL, which are considered to arise from electron beam induced damage. In the PsbO subunit, a disulfide bond between Cys19 and Csy41 was observed in the XFEL structure^10^, but it was completely cleaved in the cryo-EM structure (Fig. 3a). In the C terminus of D1 subunit, a part of Ala344, the C-terminal residue that coordinated to the Mn_4_CaO_5_ cluster, flipped out from the OEC and adopted an alternative conformation in the cryo-EM structure (Fig. 3b). These are the typical sign of damage caused by the electron beam irradiation during the acquirement of cryo-EM images.

**Fig. 3.**
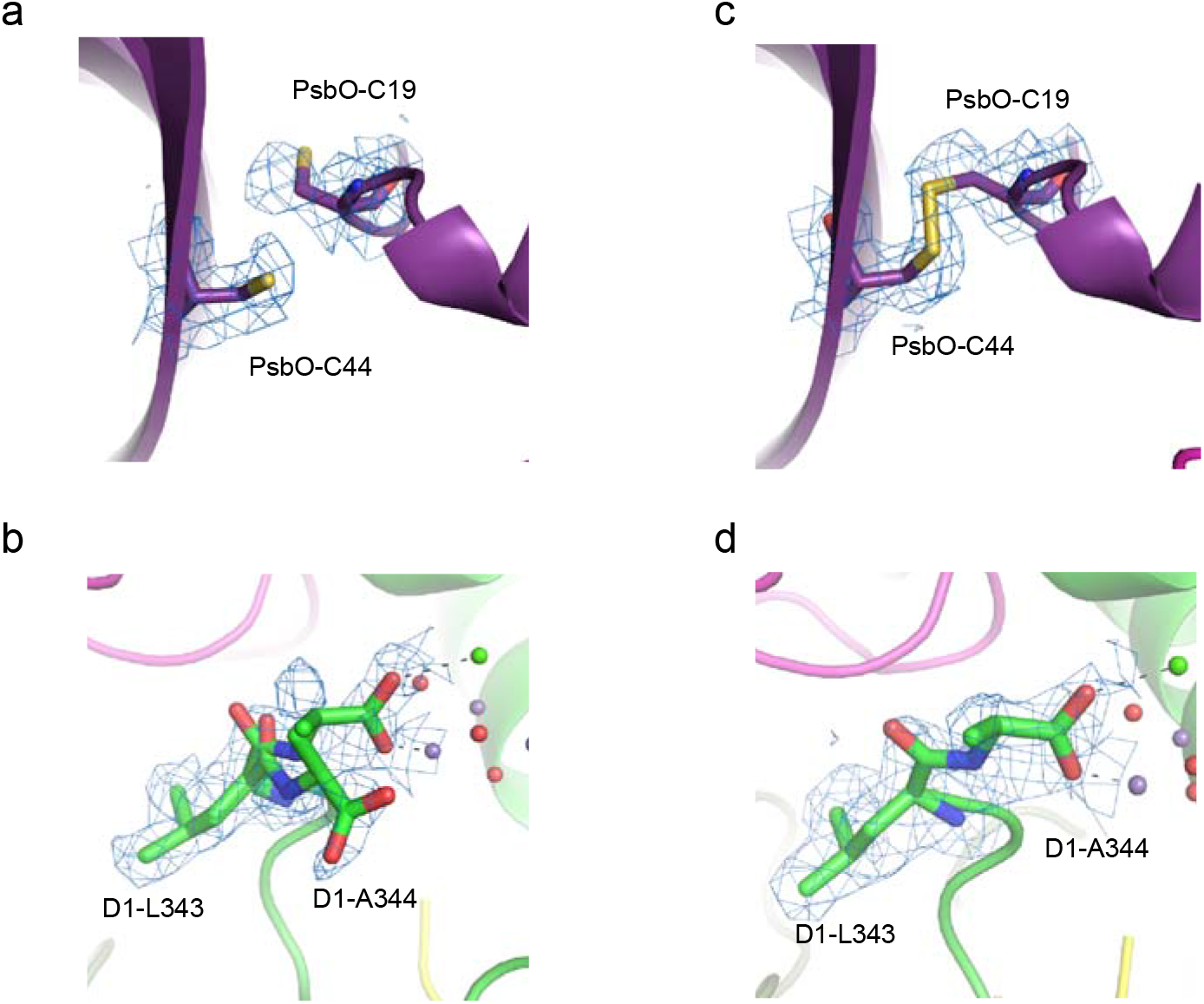
Electron beam damages in the PSII structure of the ARM-60k data set at the high-dose (83 e^−^ Å^−2^) and low-dose (3.3 e^−^ Å^−2^). **a** The broken disulfide bond in PsbO at the high-dose. **b** The alternative conformation at D1-A344 at the high dose. **c** The disulfide bond recovered in PsbO at the low-dose. **d** The single conformation of D1-A344 at the low dose. The densities were depicted at 5 σ.

In the OEC, the positions of heavy metals were confirmed clearly and were assigned based on their highest peaks in the cryo-EM map achieved at the high-dose (Fig. 4a). In addition, five oxo-oxygen atoms and four water molecules ligated to the OEC were assigned in the difference map, which were obtained by subtracting the metal densities in a diameter of 1.5 Å of that metal from the whole cryo-EM map. The overall architecture of the OEC in the cryo-EM structure is very similar to that of the SR and XFEL structures, however, distinct differences were observed in Mn–Mn and Mn–O distances (Table 3). The Mn–Mn distances calculated from the initially assigned positions based on the cryo-EM density were 2.8 Å for Mn1–Mn2, 3.5 Å for Mn1–Mn3, 5.0 Å for Mn1–Mn4, 3.1 Å for Mn2–Mn3, 5.4 Å for Mn2–Mn4, and 2.7 Å for Mn3–Mn4 (Table 3). Except the Mn1-Mn4 and Mn3-Mn4 distances, all of the Mn-Mn distances are 0.1-0.2 Å and 0.1–0.4 Å longer than those of the SR and XFEL structures, respectively^6,10^. Most of the Mn-O distances in the cryo-EM structure were also 0.1-0.5 Å and 0.1–0.7 Å longer than those in the SR and XFEL structures, respectively (Table 3)^6,10^. These differences may be caused by two factors. One is the electron beam damage, and the other one is the experimental errors in determining the positions of the individual atoms based on the cryo-EM map only. Especially, the position of oxygen atoms may not be determined precisely because the map of the oxygen atoms cannot be separated from the map of metal atoms. Thus, the OEC structure was refined with the restraints for bond distances (Mn–O and Ca–O) that were taken from the initial positions. The Mn–Mn distances refined with restraints were 2.8 Å for Mn1–Mn2, 3.4 Å for Mn1–Mn3, 5.0 Å for Mn1–Mn4, 3.1 Å for Mn2–Mn3, 5.6 Å for Mn2–Mn4, and 3.0 Å for Mn3–Mn4 (Fig. 4b and Table 3). Except the Mn1-Mn4 distance, all of the Mn-Mn distances are still 0.1-0.2 Å and 0.1–0.4 Å longer than those of the SR and XFEL structures, respectively^6,10^. Most of the Mn-O distances in the cryo-EM structure refined with the restraints were also 0.1–0.4 Å longer than those in the SR and XFEL structures (Table 3). In addition, the occupancy of the OEC atoms refined with Refmac5 were found to be lower than 1.0 (0.87). These results indicate that the OEC is reduced by electron dose exposed, leading to the elongation of the Mn-Mn and Mn-O distances and some disorder or displacement of the metal centers during the cryo-EM data acquisition. This reduced occupancy is in agreement with the previous theoretical calculation of the cryo-EM structure of higher plant PSII-LHCII supercomplex^25^, which may be the reason why a part of Ala344 flipped out and does not ligate to the OEC. The reduction of metal ions with electron doses was already observed in electron crystallography previously^26^.

**Fig. 4.**
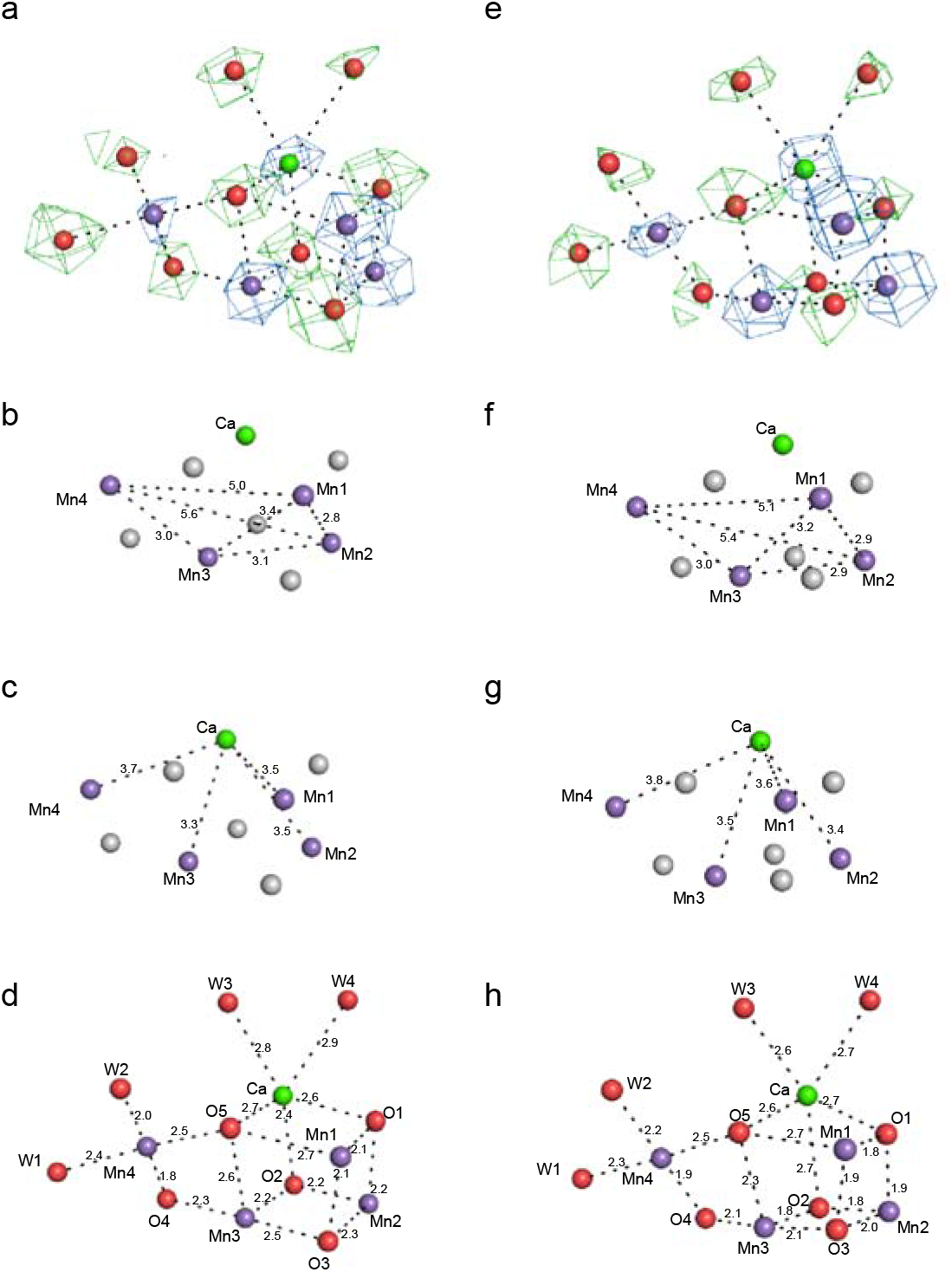
Electron beam damages in the OEC structure solved at the high-dose (83 e^−^ Å^−2^) and low-dose (3.3 e^−^ Å^−2^). **a-d:** High dose structure. **a** The cryo-EM density (blue) for manganese and calcium atoms at 17 σ and the subtracted map (green) for oxygen atoms and water molecules at 7 σ. **b** Mn– Mn distances in the OEC (in Å). **c** Mn–Ca distances in the OEC (in Å) **d** Mn–O, Ca–O, Mn–water and Ca–water distances in the OEC (in Å). **e-h:** Low dose structure. **e** The cryo-EM density (blue) for manganese and calcium atoms at 17 σ and the subtracted map (green) for oxygen atoms and water molecules at 7 σ. **f** Mn–Mn distances in the OEC (in Å). **g** Mn–Ca distances in the OEC (in Å). **h** Mn–O, Ca–O, Mn–water and Ca–water distances in the OEC (in Å).

**Table 3.** Summarization of the distances of atoms of the Mn_4_CaO_5_ cluster.

|  | High-dose | Low-dose | SR (3WU2) | XFEL (4UB6) | High-dose |  |  |  |  | Low-dose |  |  |  |  |
| --- | --- | --- | --- | --- | --- | --- | --- | --- | --- | --- | --- | --- | --- | --- |
|  | (Average) | (Average) | (Average) | (Average) | 4th | 3rd | 2nd | 1st | Initial | 4th | 3rd | 2nd | 1st | Initial |
| **Mn1-Mn2** | 2.8 | 2.9 | 2.8 | 2.7 | 2.81 | 2.83 | 2.82 | 2.82 | 2.81 | 2.88 | 2.87 | 2.89 | 2.8 | 2.95 |
| **Mn1-Mn3** | 3.4 | 3.2 | 3.3 | 3.2 | 3.35 | 3.36 | 3.35 | 3.36 | 3.46 | 3.20 | 3.20 | 3.24 | 3.18 | 3.37 |
| **Mn1-Mn4** | 5.0 | 5.1 | 5 | 5 | 5.04 | 5.02 | 5.03 | 5.03 | 4.99 | 5.10 | 5.09 | 5.08 | 5.04 | 5.32 |
| **Mn2-Mn3** | 3.1 | 2.9 | 2.9 | 2.7 | 3.08 | 3.10 | 3.09 | 3.09 | 3.06 | 2.88 | 2.88 | 2.88 | 2.85 | 2.75 |
| **Mn2-Mn4** | 5.6 | 5.4 | 5.4 | 5.2 | 5.65 | 5.65 | 5.65 | 5.63 | 5.42 | 5.47 | 5.46 | 5.42 | 5.43 | 5.40 |
| **Mn3-Mn4** | 3.0 | 3.0 | 3.0 | 2.9 | 2.97 | 2.97 | 2.98 | 2.94 | 2.67 | 3.04 | 3.03 | 3.01 | 3.00 | 3.05 |
| **Mn1-Ca** | 3.5 | 3.6 | 3.5 | 3.5 | 3.55 | 3.52 | 3.52 | 3.52 | 3.51 | 3.59 | 3.57 | 3.55 | 3.56 | 3.83 |
| **Mn2-Ca** | 3.5 | 3.4 | 3.4 | 3.3 | 3.49 | 3.45 | 3.46 | 3.46 | 3.29 | 3.44 | 3.43 | 3.41 | 3.44 | 3.35 |
| **Mn3-Ca** | 3.3 | 3.5 | 3.4 | 3.4 | 3.32 | 3.29 | 3.28 | 3.27 | 3.04 | 3.45 | 3.45 | 3.44 | 3.46 | 3.43 |
| **Mn4-Ca** | 3.7 | 3.8 | 3.8 | 3.8 | 3.73 | 3.69 | 3.70 | 3.68 | 3.57 | 3.76 | 3.76 | 3.70 | 3.80 | 3.86 |
| **Mn1-O1** | 2.1 | 1.8 | 1.9 | 1.8 | 2.11 | 2.10 | 2.08 | 2.06 | 2.34 | 1.80 | 1.76 | 1.77 | 1.78 | 2.09 |
| **Mn1-O3** | 2.1 | 1.9 | 1.8 | 1.9 | 2.09 | 2.10 | 2.10 | 2.09 | 2.29 | 1.89 | 1.90 | 1.91 | 1.92 | 1.90 |
| **Mn1-O5** | 2.7 | 2.7 | 2.6 | 2.7 | 2.72 | 2.73 | 2.74 | 2.75 | 2.64 | 2.71 | 2.72 | 2.72 | 2.72 | 2.89 |
| **Mn2-O1** | 2.2 | 1.9 | 2.1 | 1.8 | 2.18 | 2.17 | 2.17 | 2.16 | 2.19 | 1.98 | 1.95 | 1.92 | 1.86 | 2.12 |
| **Mn2-O2** | 2.2 | 1.8 | 2.1 | 1.8 | 2.24 | 2.23 | 2.2 | 2.17 | 2.45 | 1.84 | 1.83 | 1.82 | 1.80 | 2.45 |
| **Mn2-O3** | 2.3 | 2.0 | 2.1 | 2.0 | 2.28 | 2.28 | 2.27 | 2.26 | 2.36 | 2.01 | 2.01 | 2.01 | 2.00 | 1.95 |
| **Mn3-O2** | 2.2 | 1.8 | 1.9 | 1.9 | 2.18 | 2.18 | 2.16 | 2.14 | 2.04 | 1.80 | 1.78 | 1.81 | 1.86 | 2.08 |
| **Mn3-O3** | 2.5 | 2.1 | 2.1 | 2.1 | 2.48 | 2.47 | 2.44 | 2.42 | 2.48 | 2.05 | 2.06 | 2.06 | 2.07 | 2.02 |
| **Mn3-O4** | 2.3 | 2.1 | 2.1 | 1.9 | 2.30 | 2.30 | 2.28 | 2.26 | 2.38 | 2.17 | 2.11 | 2.05 | 1.96 | 2.71 |
| **Mn3-O5** | 2.6 | 2.3 | 2.4 | 2.2 | 2.58 | 2.58 | 2.56 | 2.54 | 2.58 | 2.35 | 2.32 | 2.29 | 2.23 | 2.57 |
| **Mn4-O4** | 1.8 | 1.9 | 2.1 | 2.0 | 1.80 | 1.80 | 1.79 | 1.82 | 1.96 | 1.86 | 1.89 | 1.92 | 1.97 | 2.05 |
| **Mn4-O5** | 2.5 | 2.5 | 2.5 | 2.3 | 2.49 | 2.46 | 2.45 | 2.44 | 2.59 | 2.5 | 2.48 | 2.45 | 2.40 | 2.67 |
| **Mn4-W1** | 2.4 | 2.3 | 2.2 | 2.3 | 2.38 | 2.40 | 2.39 | 2.40 | 2.54 | 2.31 | 2.32 | 2.36 | 2.32 | 2.33 |
| **Mn4-W2** | 2.0 | 2.2 | 2.2 | 2.1 | 2.05 | 2.03 | 2.03 | 2.07 | 2.11 | 2.18 | 2.18 | 2.18 | 2.15 | 1.84 |
| **Ca-O1** | 2.6 | 2.7 | 2.4 | 2.6 | 2.63 | 2.63 | 2.62 | 2.62 | 2.60 | 2.72 | 2.70 | 2.67 | 2.63 | 2.47 |
| **Ca-O2** | 2.4 | 2.7 | 2.5 | 2.7 | 2.39 | 2.39 | 2.41 | 2.43 | 2.41 | 2.67 | 2.68 | 2.69 | 2.71 | 2.73 |
| **Ca-O5** | 2.7 | 2.6 | 2.7 | 2.5 | 2.71 | 2.70 | 2.68 | 2.66 | 2.70 | 2.62 | 2.60 | 2.58 | 2.55 | 2.60 |
| **Ca-W3** | 2.8 | 2.6 | 2.4 | 2.6 | 2.76 | 2.79 | 2.78 | 2.78 | 2.80 | 2.56 | 2.56 | 2.57 | 2.53 | 2.64 |
| **Ca-W4** | 2.9 | 2.7 | 2.5 | 2.5 | 2.89 | 2.93 | 2.92 | 2.94 | 2.92 | 2.71 | 2.71 | 2.73 | 2.65 | 2.71 |

Further structural changes in the redox-active sites, including reaction center chlorophylls, electron transfer chain, proton channels and water channels, were not found except the water molecule near D2-Tyr160 (Y_D_) (Supplementary Fig. 6c). This water molecule was disordered at two positions with one being able to hydrogen bond to Y_D_ and the other one being able to hydrogen bond to D2-Arg180 in the SR structure and XFEL structures^6,10^. In the cryo-EM structure, this water molecule was ordered and connected to D2-Arg180. This may reflect the electron beam induced damage, which causes reduction of Y_D_ and broken the hydrogen-bond to Y_D_^+^.

### Reducing electron beam dosage in determining the PSII structure

In order to reduce the electron beam damage, the final cryo-EM maps were calculated from only initial several frames of each movie stack. In Supplementary Fig. 7, the inverse resolutions of reconstructions achieved from decreased electron doses for the ARM-60k data set, associated with the frame numbers, are plotted against the dose values on a logarithmic scale. Surprisingly, the electron doses from 83 e^−^ Å^−2^ down to 10 e^−^ Å^−2^ gave rise to almost the same resolution, indicating that increase in the electron beam dosage during this range does not contribute to increase in the resolution significantly. Near atomic resolution is retained at the total dose of 3.3 e^−^ Å^−2^ for the ARM-60k data set (2.08 Å), which was achieved by using the initial 2 frames of each micrograph.

An overall atomic model of the low-dose PSII was built based on the highest 2.08 Å resolution density reconstructed from the ARM-60k data set at the dose of 3.3 e^−^ Å^−2^ (Table 2). The overall architecture of the low-dose PSII is very similar to that of the high-dose PSII, with a RMSD of 0.21 Å for 5310 C_α_ atoms between the structures of high-dose and low-dose. However, in the regions where structural changes were observed due to electron beam damage, the disulfide bond between Cys19 and Csy41 of the PsbO was restored at a dose of 5 e^−^ Å^−2^, and Ala344 of the D1 subunit was returned to the single conformation to ligate to the OEC similar to those seen in the crystal structures (Fig. 3, 4 and 5, Supplementary Fig. 6 and 8). The ADP for the OEC atoms (12.8 Å^2^) were lower than that observed in the overall protein atoms of the cryo-EM structure (22.4 Å^2^), and the occupancy value of the OEC atoms refined with Refmac5 was returned to 1.0. These results indicate the reduction of the electron beam damage in the structure. However, similar to the high-dose structure, the water molecule near Y_D_ are connected to D2-Arg180 in an ordered manner and not hydrogen-bonded to Y_D_ (Supplementary Fig. 6), indicating that some electron beam damage remained.

**Fig. 5.**
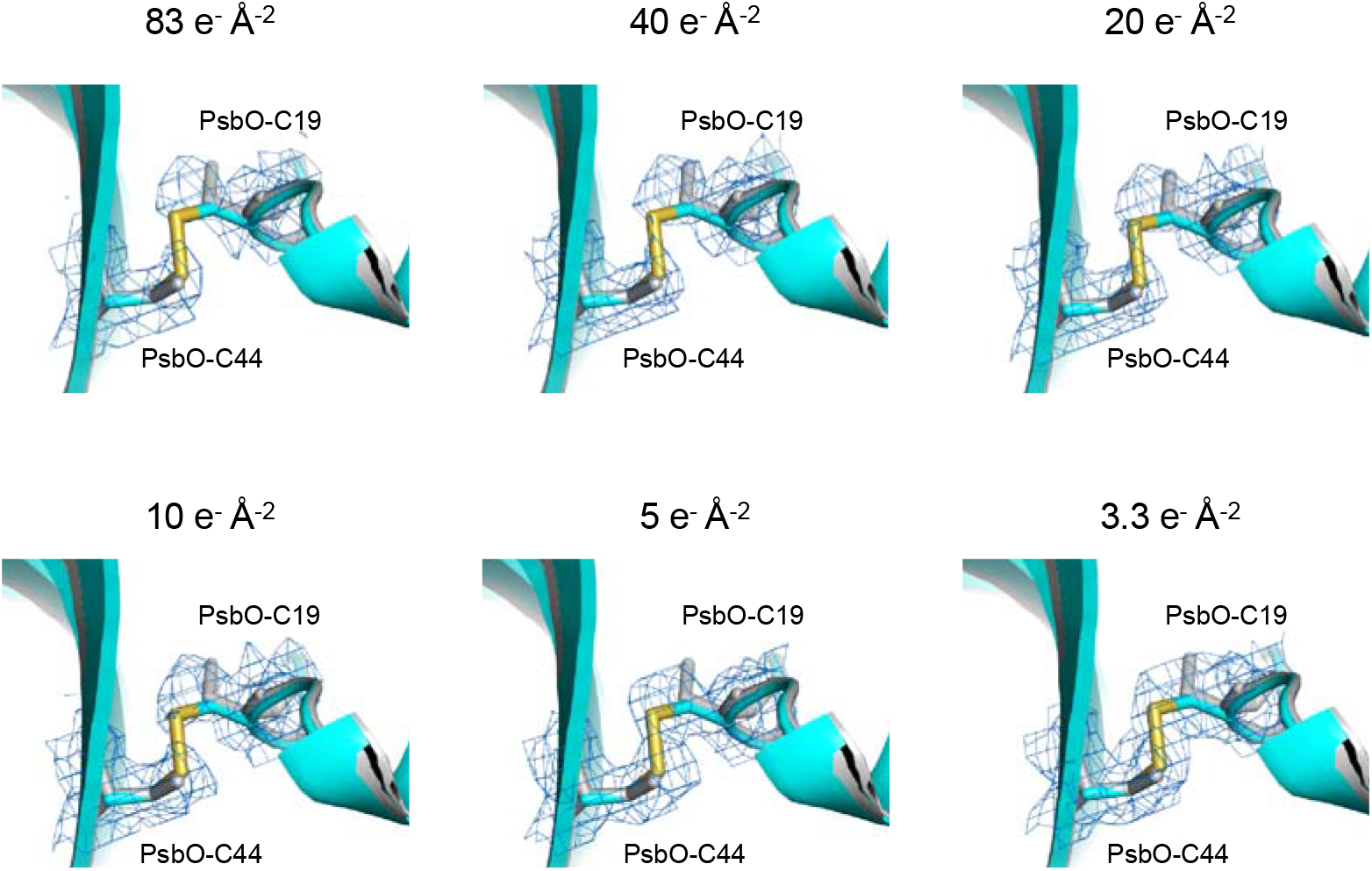
Changes of the cryo-EM map in the region of the disulfide bond in PsaO with changes of the electron beam dose. The cryo-EM maps for each electron dose are displayed as a blue mesh at 4 σ and the corresponding models for low-dose (colored) and high-dose (gray) are shown as sticks.

The Mn–Mn distances calculated from the initially assigned positions based on the cryo-EM density were 3.0 Å for Mn1–Mn2, 3.4 Å for Mn1–Mn3, 5.3 Å for Mn1–Mn4, 2.8 Å for Mn2–Mn3, 5.4 Å for Mn2–Mn4, and 3.1 Å for Mn3–Mn4 (Table 3). All of these Mn-Mn distances are 0.1–0.3 Å longer than those of the SR and XFEL structures^6,10^. Most of the Mn-O distances in the cryo-EM structure were also 0.1–0.6 Å and 0.1–0.8 Å longer than those in the SR and XFEL structures, respectively (Table 3). As is done with the high-dose structure, we refined the OEC structure with the restraints for bond distances of Mn–O and Ca–O that were taken from the initial positions. The Mn–Mn distances in the OEC refined with the restraints were 2.9 Å for Mn1–Mn2, 3.2 Å for Mn1–Mn3, 5.1 Å for Mn1–Mn4, 2.9 Å for Mn2–Mn3, 5.4 Å for Mn2–Mn4, and 3.0 Å for Mn3–Mn4 (Fig. 4f, Table 3). Most of these distances are shorter than those of the high-dose structure and close to those of the XFEL structure, although most of them are still longer than the SR and XFEL structures by 0.1 Å and 0.1-0.2 Å, respectively (Table 3). The Mn-O distances after refinement with the restraints also became close to the XFEL structure, indicating the necessity of refinement with restraints. However, some of the Mn-O distances were still longer or deviated from those found in the XFEL structure, which may be caused by the electron beam damage remained and/or coordinate errors in the cryo-EM analysis at the current resolution.

## Discussion

In recent years, the resolution of single-particle cryo-EM has improved to atomic resolutions without crystallization, and it has been reported that biological samples are damaged by electron beams. In this study we elucidated the structure of the PSII at 1.95 Å resolution in solution by cryo-EM, which is similar to the SR or XFEL structure in the crystalline state except the PsbY subunit, which was visible in the cryo-EM structure but absent or partially visible in the SR and XFEL structures^6,10^. This indicates that the structure solved by cryo-EM more closely represents the physiological state.

Despite the total electron dose of 83 e^−^ Å^−2^ which is commonly used in the acquisition of cryo-EM images, radiation damages are found in regions susceptible to redox changes, i.e. the disulfide bond and the redox-active metals, whereas the overall structure is very similar to those of the SR and XFEL structures^6,10^. The exposure of the sample to a flux of electrons is conveniently expressed in terms of electrons per Å^2^ of specimen surface area (e^−^ Å^−2^), which is converted to the SI unit for the absorbed ionizing radiation, the Gray (Gy, with 1 Gy = 1J kg^−1^), by a factor of □3.7 ^27^. Thus, the total electron dose of 83 e^−^ Å^−2^ is equal to the absorption of 307 MGy, which greatly exceeds the Henderson limit (20 MGy) that is the X-ray dose that a cryo-cooled crystal can absorb before the diffraction pattern decays to half of its original intensity^28^. Nevertheless, our cryo-EM structure is almost the same to the SR and XFEL structures in the redox-active sites, including reaction center chlorophylls, electron transfer chain, proton channels and water channels, indicating that the radiation damage does not affect the structure significantly. This is considered to be the result of successful dose-weighted correction in the Bayesian polishing step. However, in the PsbO subunit, disulfide bond between Cys19 and Csy41 were completely breaking (Fig. 3) in the cryo-EM structure. In the OEC structure, the Mn-Mn were 0.1–0.4 Å longer than those in the XFEL structure, and most of the Mn-O distances were also significantly longer. In addition, the occupancy of the OEC atoms were lower than 1.0 (0.87), resulting in a multiple conformation of the C-terminal residue of D1, where a part of Ala344 flipped to a direction that does not ligate to the OEC.

We examined whether radiation damage could be reduced by reducing the total number of stacked movie frames used in the structural analysis. In the electron doses analyzed, the reconstructed map from summing the initial two frames of each micrograph (3 e^−^ Å^−2^, 11.1 MGy) gave rise to almost similar resolutions to that of high-dose data set (83 e^−^ Å^−2^, 307 MGy) (Supplementry Fig.7). The reconstructed map from the ARM-60k data set at the dose of 3.3 e^−^ Å^−2^ (11.1 MGy) gave a resolution of 2.08 Å. This structure was compared with that obtained with the high-dose, and it was found that the disulfide bond in the PsbO was recovered, and Ala344 of the D1 subunit was returned to the single conformation similar to the SR and XFEL structures (Fig. 3 and 5). This indicates a significant reduction of the electron beam damage to the structure. In the structure of OEC, most of the Mn-Mn and Mn-O distances became shorter than those observed in the high-dose structure before refinement. However, some of them are still significantly deviated from those of the XFEL structure (Table 3), which become closer after the refinement with restraints starting from the initial structure. Thus, it is advisable to refine the cryo-EM structure with restraints imposed for compounds such as the Mn_4_CaO_5_ cluster.

After refinement, most of the Mn-Mn and Mn-O distances of the low-dose structure are similar to those observed in the XFEL structure. However, some of the distances are still longer than or deviated from those found in the damage free XFEL structure^10^. This may be caused by two reasons; one is some electron beam damage remained, and the other one is coordinate errors existed in the cryo-EM structure. It has been reported that about 80% of Mn of OEC in solution is reduced to divalent cations by an X-ray dose of 5 MGy^7^. Even though it is estimated that about 90% of Mn of OEC is Mn(II) at an electron beam dose of 11.1 MGy used for the low-dose structure, the structure of the OEC retained an occupancy of 1.0. This may be contributed by the stability of the structure of OEC, as the metal ions of OEC are liganded by seven amino acid residues (D1-D170, D1-E189, D1-H332, D1-E333, D1-D342, D1-A344, and CP43-E354). However, the longer distances observed in some of the metal-oxygen distances of OEC even in the low-dose structure indicated the existence of electron beam damage. Tanaka et al. has reported that, using SR, a dose of 0.1 MGy is necessary to achieve a structure similar to that of the XFEL structure^29^. Thus, in order to achieve a damage free structure, the electron beam dose needs to be further reduced. Fortunately, our data indicated that the resolution depends on the particles used, and by using more images and particles, it will be possible to lower the electron beam dose and achieve the structure at a higher resolution.

Coordinate errors in the cryo-EM structure may be caused by the ambiguities in the orientations of particles and their averaging, as well as the subsequent structural analysis procedures. Structural analysis by cryo-EM at a higher resolution should eliminate such errors, and gives rise to a more accurate structure. It is also expected that improvements in the averaging and structural analysis algorithms of the cryo-EM data may improve the accuracy of the structures at the same resolutions.

In summary, we show that the electron dose commonly used in cryo-EM is damaging to protein samples. However, the damaged area was limited to redox-sensitive part. Our results suggest that it is possible to obtain a structure with less damage and high resolution by reducing the total dose and increasing the number of particles. This study will serve as a good indicator for determining damage-less cryo-EM structures of PSII and all biological samples, especially redox-active metalloproteins.

## Methods

### Growth of cells and purification of PSII

Cells of *Thermosynechococcus vulcanus* (*T. vulcanus*) were grown in four 10 L bottles at 50°C. PSII with a high oxygen evolving activity was purified from *T. vulcanus* as described previously^30–32^ and suspended with a buffer containing 20 mM MES-NaOH (pH 6.0), 0.04% β-dodecyl-D-maltopyranoside and 5% glycerol. For the Titan-96k and ARM-60k data collection, glycerol in the buffer was removed by polyethylene glycol (PEG) precipitation and the resultant PSII was re-suspended in a buffer containing 20 mM MES-NaOH (pH 6.0), 20 mM NaCl, 3 mM CaCl_2_, 0.04% *β*-dodecyl-D-maltopyranoside.

### Cryo-EM data collection

For cryo-EM experiments, 3-μL aliquots of the PSII sample at each condition (shown in Table 1) were applied to Quantifoil R1.2/1.3, Mo 300 mesh or Cu 200 mesh grids. The grids were incubated for 10 s in an FEI Vitrobot Mark IV at 4°C and 100% humidity. The grids were immediately plunged into liquid ethane cooled by liquid nitrogen and then transferred into the Titan Krios (Thermo Fischer Scientific) equipped with a field emission gun, a Cs corrector (CEOS GmbH), and a direct electron detection camera (Falcon 3EC, Thermo Fischer Scientific), or CRYO ARM 300 (JEOL) equipped with a cold-field emission gun and a direct electron detection camera (Gatan K2 summit, Gatan Inc). These microscopes were operated at 300 kV and a nominal magnification of × 75,000 (Titan-75k), × 96,000 (Titan-96k) for Titan Krios and × 60,000 (ARM-60k) for CRYO ARM 300. Images were recorded using the Falcon 3EC in linear mode or Gatan K2 summit in counting mode. Micrographs were recorded with a pixel size of 0.870 Å, 0.678 Å and 0.822 Å at a dose rate of 40 electrons Å^−2^ sec^−1^, 40 electrons Å^−2^ sec^−1^ and 83 electrons Å^−2^ sec^−1^ for Titan-75k, Titan-96k and ARM-60k, respectively. The nominal defocus range were −1.0 to −2.0 μm, −1.0 to −2.5 μm, and −0.8 to −1.6 μm for Titan-75k, Titan-96k and ARM-60k, respectively. Each exposure was conducted for 45.11 s, 26.64 s and 10.00 s, and were dose-fractionated into 78, 39 and 50 movie frames for Titan-75k, Titan-96k and ARM-60k, respectively. We acquired 2,084, 4,237 and 2,160 images for the data sets of Titan-75k, Titan-96k and ARM-60k, respectively.

### Cryo-EM image processing

Movie frames were aligned and summed using the MotionCor2 software^33^ to obtain a final dose weighted image. Estimation of the contrast transfer function (CTF) was performed using the CTFFIND4 program^34^. All of the following processes were performed using RELION3.0 ^35^. For structural analyses of the Titan-75k data set, 354,233 particles were automatically picked from 2,084 micrographs and then were used for reference-free 2D classification. Then, 309,028 particles were selected from the good 2D classes and subjected to 3D classification with a C2 symmetry. The 1.9 Å PSII structure from *T. vulcanus* (PDB: 3WU2)^6^ was employed for the initial model for the first 3D classification with 60-Å low-pass filter. As shown in the Supplementary Fig. 1 and 2, the PSII structure was reconstructed from 90,897 particles at an overall resolution of 2.22 Å. For structural analyses of the Titan-96k data set, 612,287 particles were automatically picked from 4,237 micrographs and then used for reference-free 2D classification. Then, 566,145 particles were selected from the good 2D classes and subjected to 3D classification with a C2 symmetry. The 2.22-Å map from Titan-75k data was employed for the initial model for the first 3D classification with a 60-Å low-pass filter. As shown in the Supplementary Fig. 1 and 2, the PSII structure was reconstructed from 203,912 particles at an overall resolution of 2.20 Å. For structural analyses of the ARM-60k data set, 481,946 particles were automatically picked from 2,160 micrographs and used for reference-free 2D classification. Then, 481,927 particles were selected from the good 2D classes and subjected to 3D classification with a C2 symmetry. The 2.22-Å map from Titan-75k data was employed for the initial model for the first 3D classification with a 60-Å low-pass filter. The PSII structure was reconstructed from 174,099 particles at an overall resolution of 1.95 Å (Supplementary Fig. 3 and 4). For the low-dose maps, the summing number of movie frames were decreased in the final step of Bayesian polishing and used for reconstruction without refinement of particle positions and orientations, using RELION^35^ with the command line option “relion_reconstruct” and then post-processed in RELION^35^. All of the resolution was estimated by the gold-standard Fourier shell correlation (FSC) curve with a cut-off value of 0.143 (Supplementary Fig. 2 and 4)^36^. The local resolution was estimated using RELION^35^.

### B-factor estimation

For the B-factor plot, the total set of all particles from the final refinement was randomly resampled into smaller subsets. These subsets were subjected to 3D auto-refinement and the resulting orientations were used to calculate reconstructions for each of the two random halves used in the auto-refinement. The squared values of the resulted, estimated resolutions were then plotted against the natural logarithm of the number of particles in the subset, and B-factors were calculated from the slope of the straight line best fitted with the points in the plot (Fig.1).

### Model building and refinement

The 1.95-Å and 2.08-Å cryo-EM maps were used for model building of the high-dose and low-dose PSII structures, respectively. First, the crystal structure of *T. vulcanus* PSII (PDB: 3WU2) was manually fitted into each cryo-EM map using UCSF Chimera^37^, and then the structures were inspected and adjusted individually with COOT^38^. The structures of high-dose PSII and low-dose PSII were then refined with phenix.real_space_refine^39^ and Refmac5^40^ with geometric restraints for the protein–cofactor coordination. The positions of four manganese atoms and one calcium atom were clearly visible in the cryo-EM map (Fig. 2 and 4). The positions of the five oxo-oxygen atoms and four water molecules ligated to the OEC were less clear, and they were identified by the difference map in which, the maps of metal ions with a diameter of 1.5 Å from that metal ion were subtracted from the whole cryo-EM map, after placement of the manganese and calcium atoms (Fig. 2 and 4). The initial positions of metal and oxygen atoms were assigned based on the highest peaks in the cryo-EM maps. This was taken as the initial structure. Subsequently, we performed the structural refinement with loose restraints (0.1 σ) for bond distances (Mn–O and Ca–O) that were taken from the initial position. Then the refinement was performed with tighter restraints (0.05 σ) for bond distances successively using the modified ‘new’ library for the bond distances. This geometry optimization procedure was repeated several times until the bond distances converged. However, the distance of Mn4-O4 in the high-dose structure, and the distances of Mn1-O1 and Mn3-O2 in the low-dose structure, were fixed to 1.8 Å, because these distances were too close and could not be refined. The averages of the distances of Mn–Mn, Mn–Ca, Mn–O and Mn–ligand were calculated from each PSIIs in the final four refinement steps and are listed in Table 3. The final models were further validated with Q-score^41^, MolProbity^42^ and EMringer^43^. The statistics for all data collection and structure refinement are summarized in Table 1 and 2. All structural figures are made by Pymol^44^ or UCSF ChimeraX^45^.

### Difference map analysis between low-dose PSII and high-dose PSII

A difference map was calculated by subtracting high-dose map from the low-dose map, i.e. (low-dose PSII) minus (high-dose PSII). The rotational and translational matrix was calculated based on the refined atomic coordinates using lsqkab in CCP4^46^. A map of low-dose PSII was superposed with a map of high-dose PSII which were applied by a low-pass filter and adjusted to 2.08 Å resolution, with calculated rotational and translational matrix using maprot in CCP4^47^. The high-dose PSII map and the low-dose PSII map was normalized based on the ratio of the root mean square map density value and then the difference maps were calculated using UCSF Chimera^37^ with the command line option “vop subtract” (Supplementary Fig. 8).

### Data availability

Atomic coordinates and cyro-EM maps for the reported structure of PSII determined from the high-dose data set of ARM-60K and low-dose data set of ARM-60k were deposited in the Protein Data Bank under an accession codes 7D1T and 7D1U, respectively, and in the Electron Microscopy Data Bank under the accession codes EMD-30547, EMD-30548, respectively. The cryo-EM maps of the Titan-75k data set and Titan-96k data set were deposited in the Electron Microscopy Data Bank under the accession codes EMD-30549 and EMD-30550, respectively.

## Acknowledgements

This work was supported by JSPS KAKENHI No. JP20H02914 (K.K.), JP19K22396, JP20H03194 (F.A.), JP20H05087 (N.M.), JP17H06434 (J.-R.S.), JST PRESTO No. JPMJPR16P1 (F.A.), the Platform Project for Supporting Drug Discovery and Life Science Research (Basis for Supporting Innovative Drug Discovery, Life Science Research (BINDS)) of AMED No. JP18am0101072j002 (N.M.), and the Cyclic Innovation for Clinical Empowerment (CiCLE) from AMED (K.Y.).

## Author Contributions

J.-R.S. and K. Y. conceived the project; Y.N. and F.A. purified the PSII; N.M. and T.H. collected cryo-EM images; N. M., K.K. and T.H. processed the EM data. K.K. built the structure model and refined the final models; K.K. analyzed the structure; and K.K., T.H., N.M. and J.-R.S. wrote the paper, and all of the authors joined the discussion of the results.

## Notes

### Competing Interest Statement

The authors have declared no competing interest.

