## Supplementary materials for "High-resolution cryo-EM structure of photosystem II: Effects of electron beam damage"

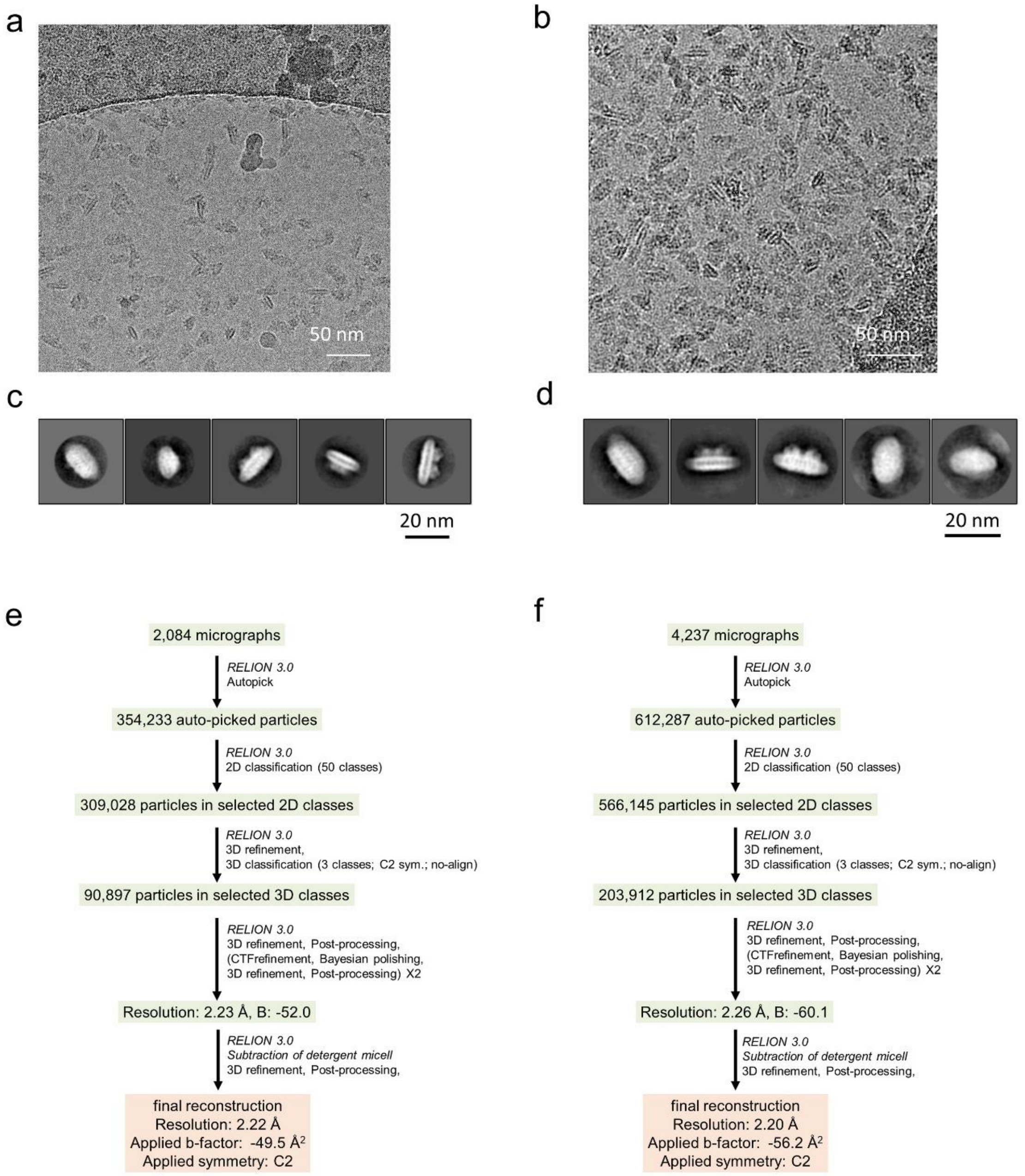


**Supplementary Fig. 1 Cryo-EM data collection and processing of PSII from Titan-75k and Titan-96k**. **a-b** A representative cryo-EM micrograph of the PSII by Titan-75k (**a**) and Titan-96k (**b**). **c-d** Representative 2D classes of the PSII particles from micrographs taken by Titan-75k (**c**) and Titan-96k (**d**). **e-f** A schematic flowchart showing the classification scheme for the PSII from Titan-75k (**e**) and Titan-96k (**f**).

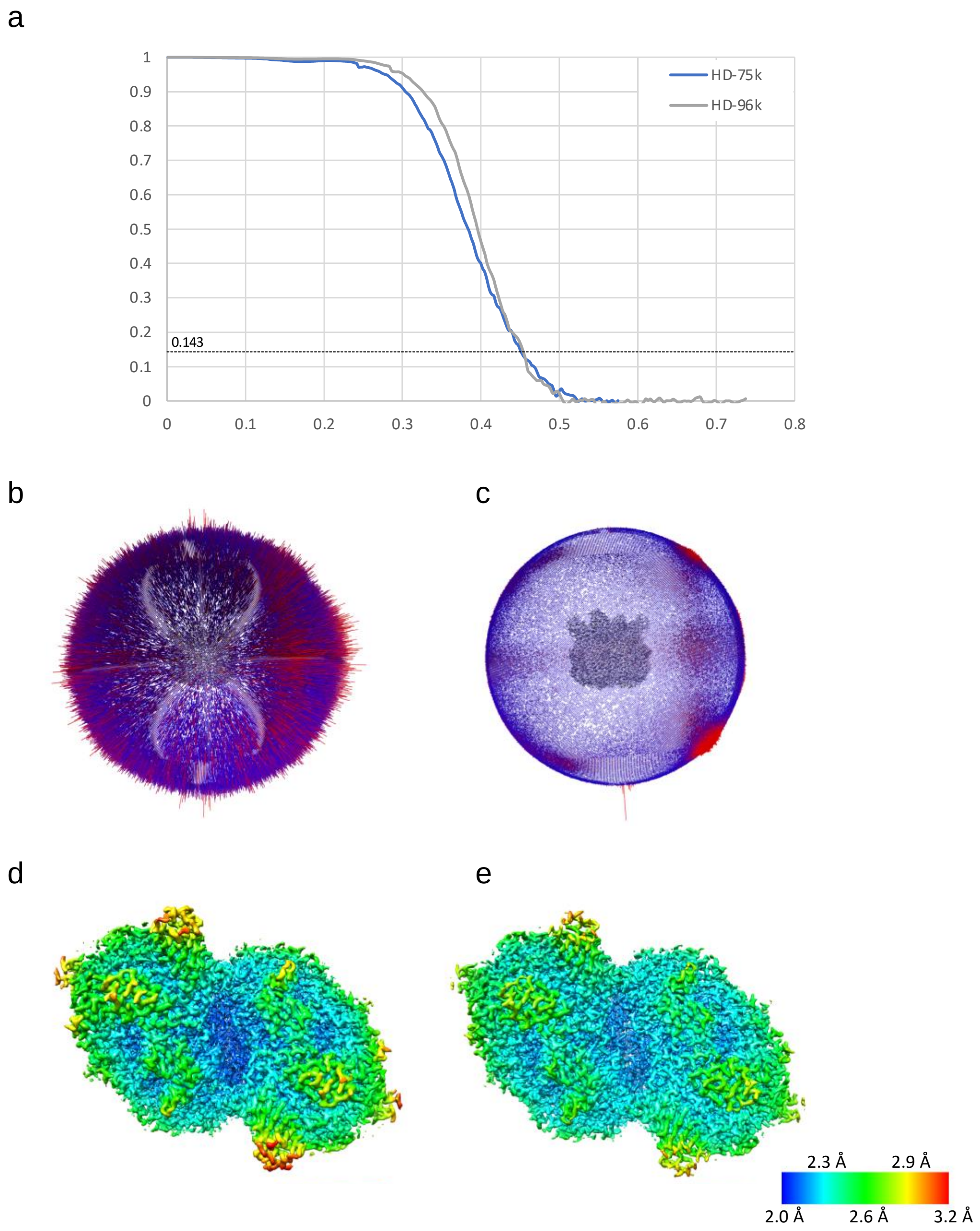


**Supplementary Fig. 2 Evaluation of the resolution of the cryo-EM map of PSII for the Titan-75k and Titan-96k data sets. a** FSC curves of the PSII for high-dose Titan-75k (blue) and high-dose Titan-96k (gray) data sets, calculated between independently refined half maps used for the structure reconstructions. **b-c** Angular distribution of the particles used for reconstruction of the PSII from Titan-75k (**b**) and Titan-96k (**c**). Each cylinder represents one view and the height of the cylinder is proportional to the number of particles for that view. **d-e** Local resolution maps of the PSII from Titan-75k (**d**) and Titan-96k (**e**).

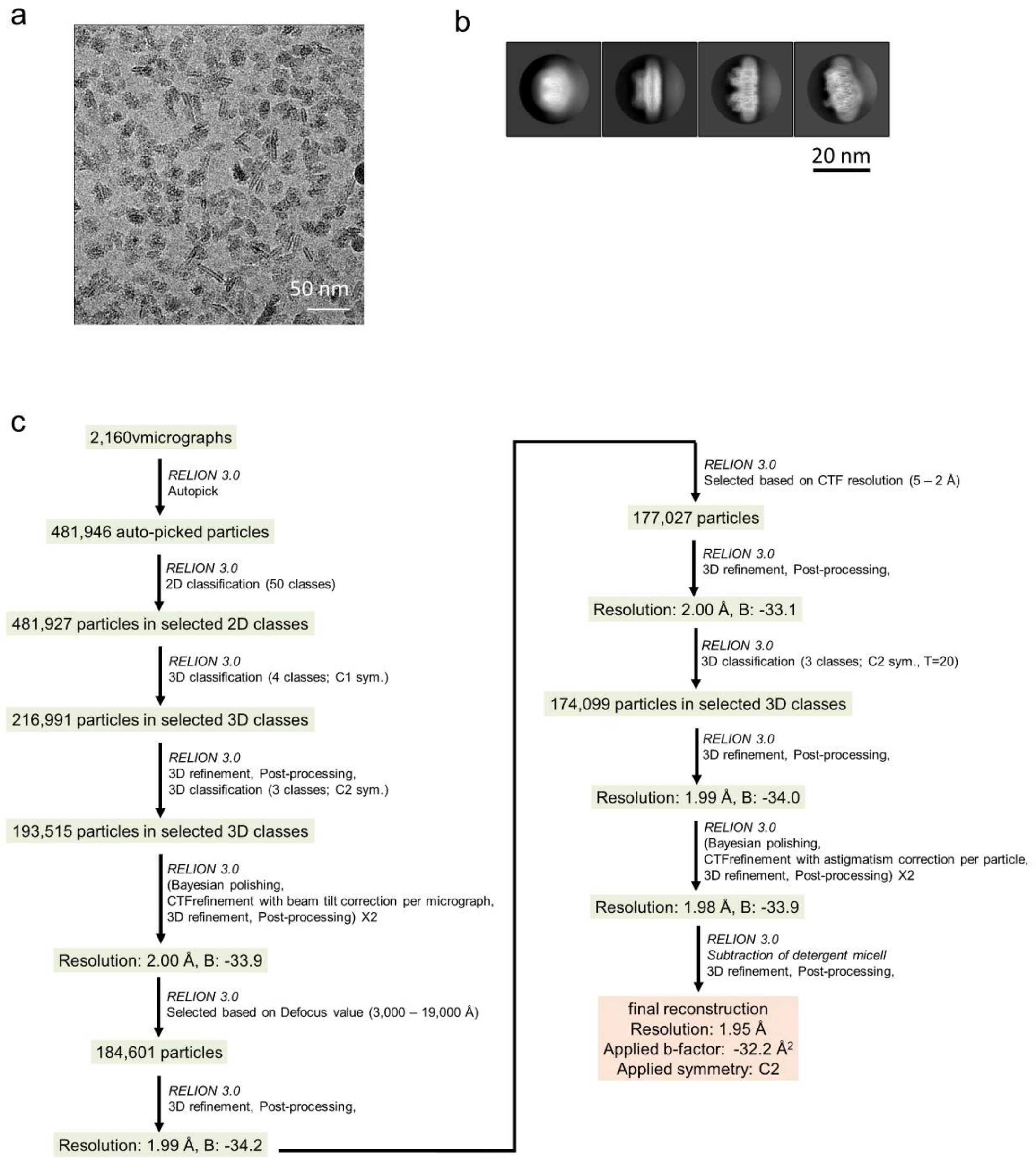


**Supplementary Fig. 3 Cryo-EM data collection and processing of PSII by ARM-60k.** **a** A representative cryo-EM micrograph of the PSII. **b** Representative 2D classes of the PSII particles. **c** A schematic flowchart showing the classification scheme for the PSII.

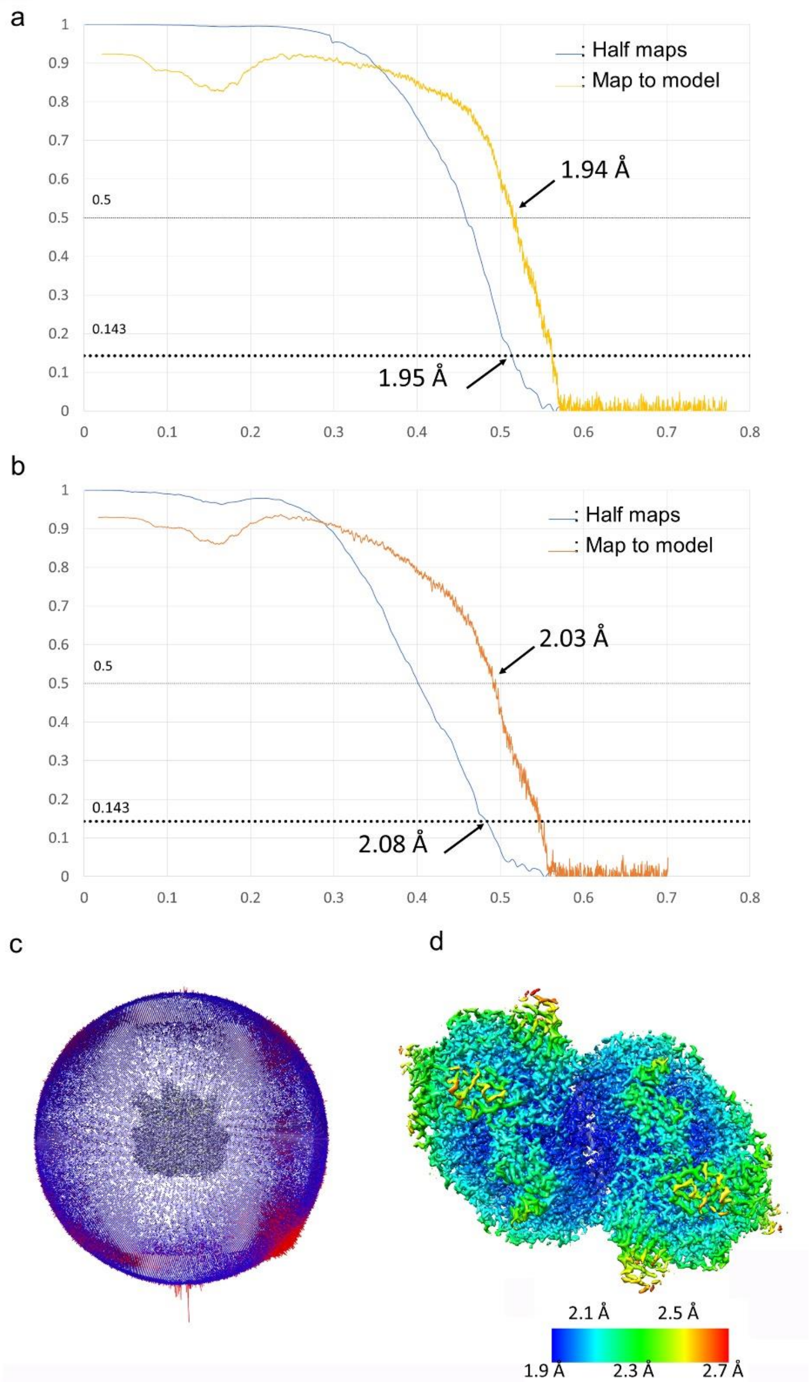


**Supplementary Fig. 4 Evaluation of the resolution of the cryo-EM map of PSII for ARM-60k. a** FSC curves of the high-dose PSII for independently refined half-maps (half-maps) and full map vs. model (map to model). **b** FSC curves of the low-dose PSII for independently refined half-maps (half-maps) and full map vs. model (map to model). **c** Angular distribution of the particles used for reconstruction of the high-dose PSII. Each cylinder represents one view and the height of the cylinder is proportional to the number of particles for that view. **d** Local resolution maps of the high-dose PSII.

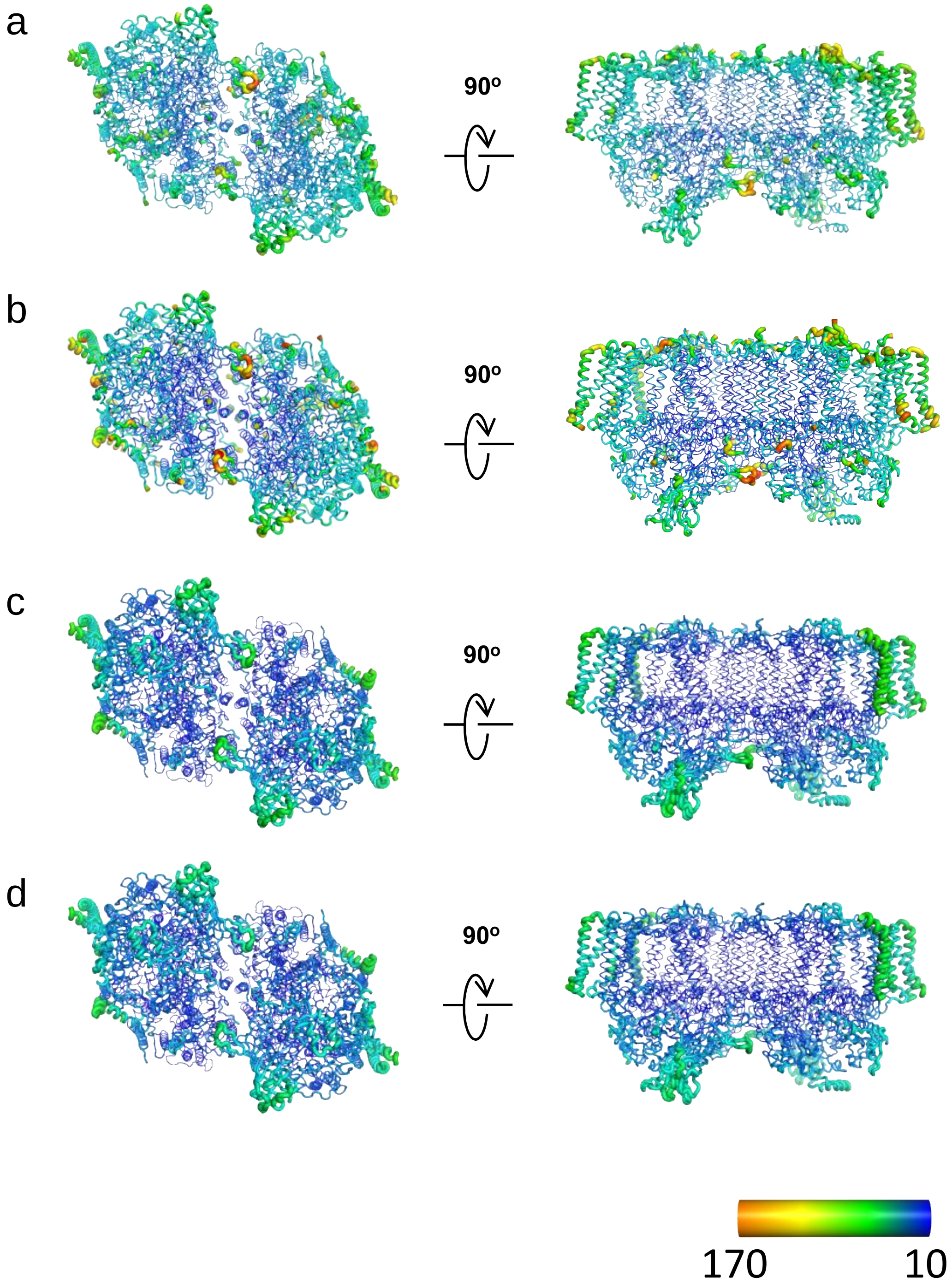


**Supplementary Fig. 5 Atomic displacement parameter (ADP) for crystal structure and cryo-EM structures. a-c** Refined ADPs for the SR structure (3WU2) (**a**), XFEL structure (4UB6) (**b**), high-dose of ARM-60k (**c**) and low-dose of ARM-60k data sets (**d**) were shown as heat maps.

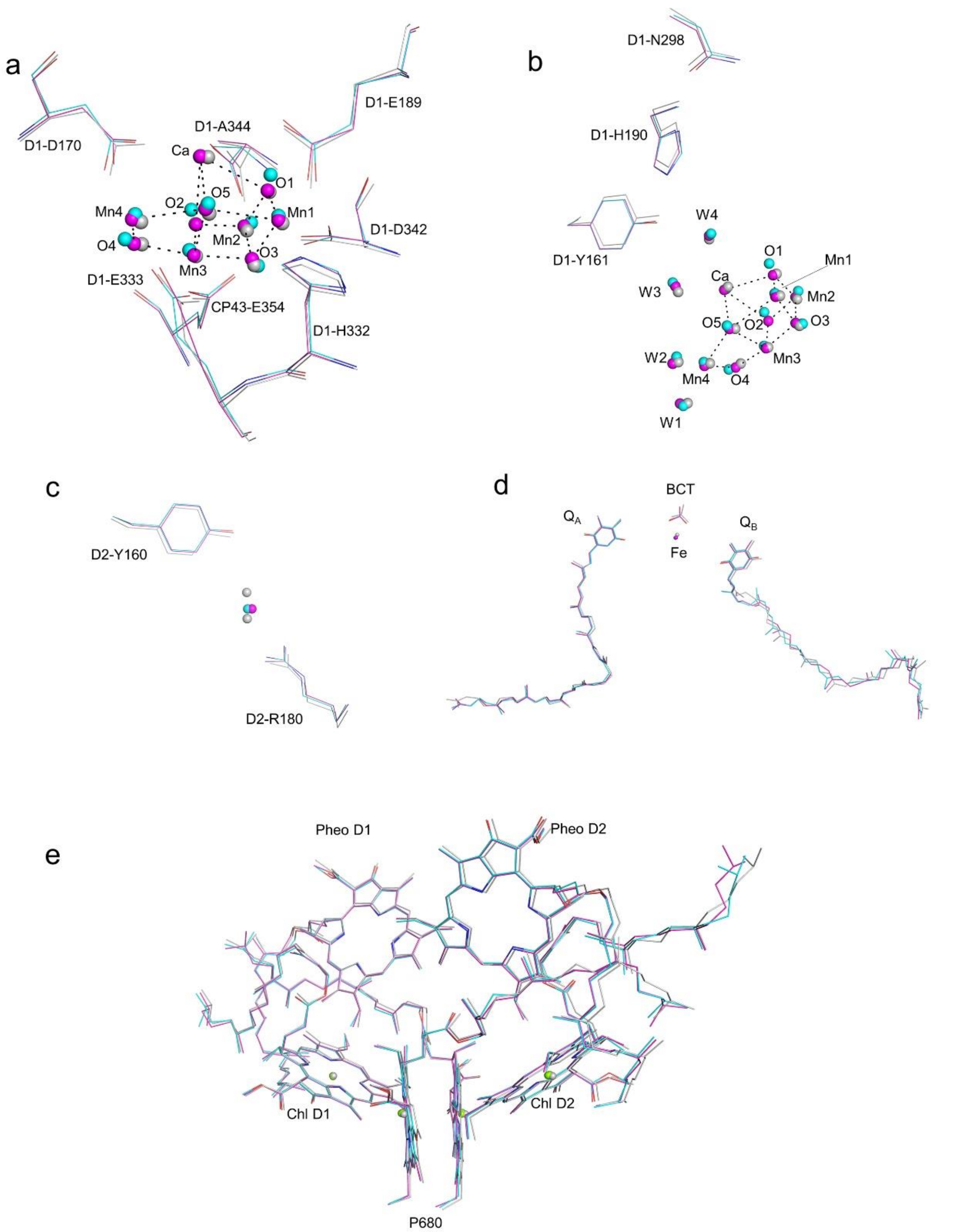


**Supplementary Fig. 6 Comparison of high-dose structure (cyan), low-dose structure (magenta) solved by cryo-EM and the XFEL structure (4UB6) (gray). a** The $Mn_4CaO_5$ cluster and its ligand environment. **b** Hydrogen bond around $Y_Z$ (D1-Tyr 161). **c** Hydrogen bond around $Y_D$ (D2-Tyr 160). **d** The $Q_A$-bicarbonate (BCT)-$Q_B$ site. **e** The P680 site.

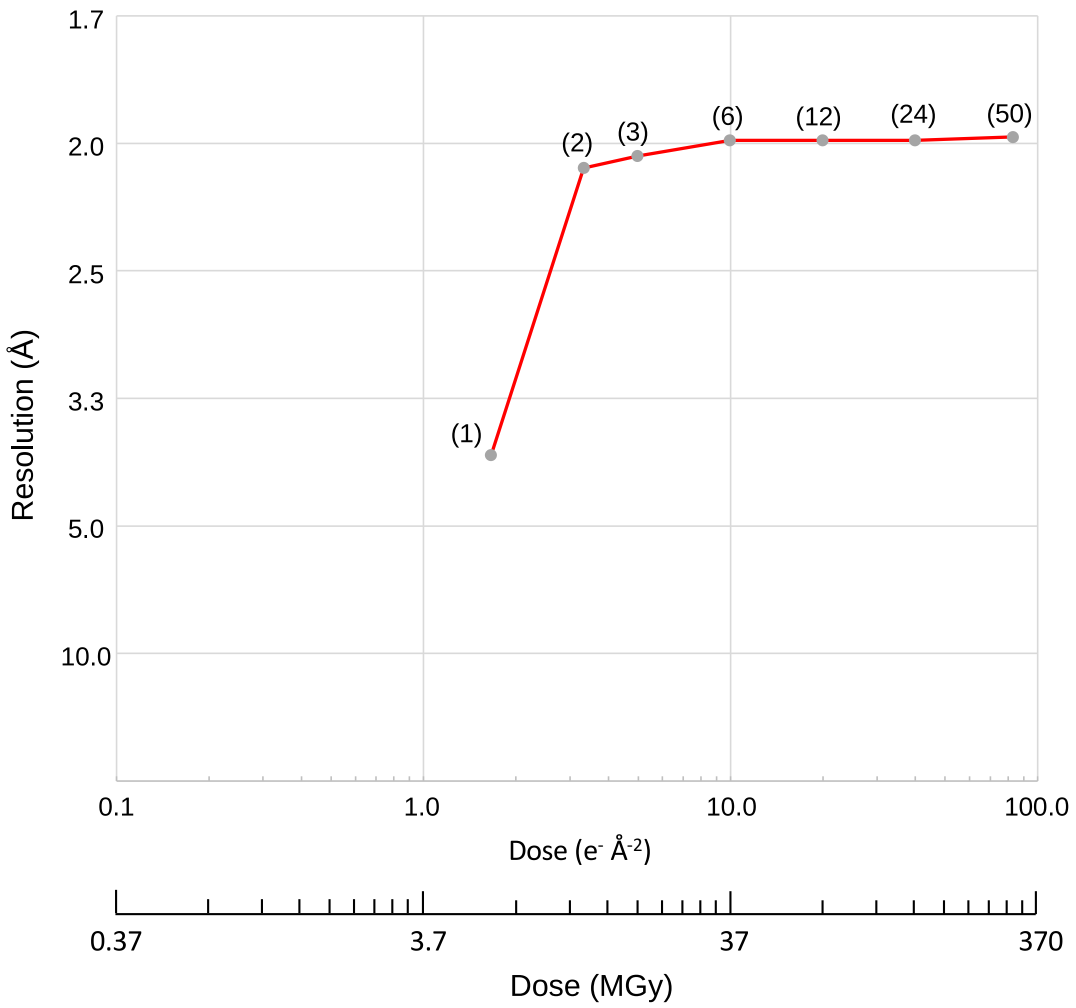


**Supplementary Fig. 7 Plot of resolutions achieved against the electron doses.** Resolutions achieved by reconstructions using each reduced frames of the total movies for data set acquired with ARM-60k. The parentheses in the graph indicate the number of frames.

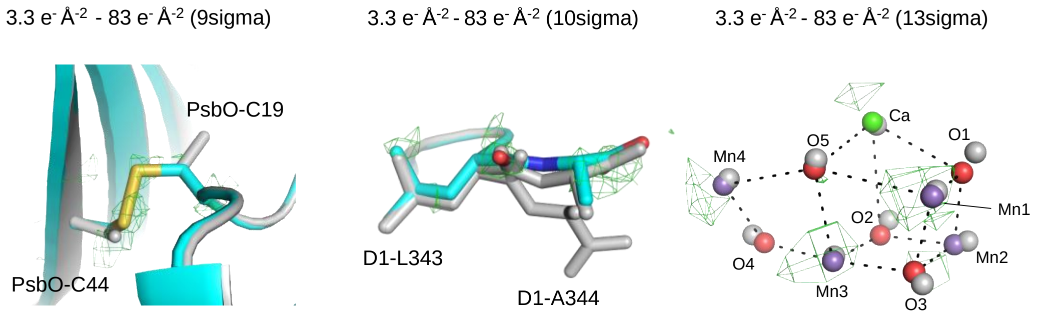


**Supplementary Fig. 8 Difference maps at the damaged parts.** The difference maps (low-dose map minus high-dose map) for each damaged area is displayed as a green mesh and the corresponding models for low-dose (colored) and high-dose (gray) are shown as sticks (see Methods).
